# SGLT2 inhibitors therapy protects glucotoxicity-induced β-cell failure in a mouse model of human K_ATP_-induced diabetes trough mitigation of oxidative and ER stress

**DOI:** 10.1101/2021.09.17.460837

**Authors:** Zeenat A. Shyr, Zihan Yan, Alessandro Ustione, Erin M. Egan, Maria S. Remedi

## Abstract

Progressive loss of pancreatic β-cell functional mass and anti-diabetic drug responsivity are classic findings in diabetes, frequently attributed to compensatory insulin hypersecretion and β-cell exhaustion. However, loss of β-cell mass and identity still occurs in mouse models of human K_ATP_-gain-of-function induced Neonatal Diabetes Mellitus (NDM), in the absence of insulin secretion. Here we studied the mechanisms underlying and temporal progression of glucotoxicity-induced loss of functional β-cell mass in NDM mice, and the effects of sodium-glucose transporter 2 inhibitors (SGLT2i) therapy. Upon tamoxifen induction of transgene expression, NDM mice developed severe diabetes followed by an unexpected loss of insulin content, decreased proinsulin processing and proinsulin accumulation at 2-weeks of diabetes. This was accompanied by a marked increase in β-cell oxidative and ER stress, without changes in islet cell identity. Strikingly, early treatment with the SGLT2 inhibitor dapagliflozin restored insulin content, decreased proinsulin:insulin ratio and reduced oxidative and ER stress. However, despite reduction of blood glucose, dapagliflozin therapy was ineffective in restoring β-cell function in NDM mice when tit was initiated at >40 days of diabetes, when loss of β-cell mass and identity had already occurred. These results have important clinical implications as they demonstrate that: *i)* hyperglycemia *per se*, and not insulin hypersecretion, drives β-cell failure in diabetes, *ii)* recovery of β-cell function by SGLT2 inhibitors is through reduction of oxidative and ER stress, *iii)* SGLT2 inhibitors revert/prevent β-cell failure when used in early stages of diabetes, but not when loss of β-cell mass/identity already occurred, *iv)* common execution pathways underlie loss and recovery of β-cell function in different forms of diabetes.

## 1. Introduction

Reduced pancreatic β-cell function and mass contribute to both type 1 diabetes (T1D) and type 2 diabetes (T2D) [1]. Reduction of β-cell function is considered an early event, while loss of β-cell mass occurs closer to clinical manifestation [2–5]. Chronic high glucose induce β-cell membrane hyperexcitability, persistently elevated intracellular calcium concentration and insulin hypersecretion, all critically contributing to loss of β-cell mass in diabetes [6–8]. However, loss of β-cell mass still occurs in K_ATP_-gain-of-function (K_ATP_-GOF) mouse model of human neonatal diabetes mellitus (NDM), in which all of these factors are absent due to K_ATP_ overactivity in pancreatic β-cells [9–13]. Chronic hyperglycemia can also induce β-cell overstimulation, oxidative and endoplasmic reticulum (ER) stress leading to β-cell exhaustion, loss of β-cell mass and identity [4, 7, 14–16]. However, most of the studies have been performed in human pancreases from T2D individuals and in animal models of T2D and obesity, and little is known about mechanisms underlying and temporal progression of loss of functional β-cell mass in monogenic diabetes, in the absence of compensatory increase in insulin secretion.

Loss of functional β-cell mass in diabetes seems to be independent of the antidiabetic therapy, with most antidiabetic agents initially effective as monotherapy, but failing to reduce blood glucose over time [17–19], requiring add-on combinational therapies. Sodium glucose transporter 2 inhibitors (SGLT2i) are a new class of antidiabetic drugs that inhibit glucose reabsorption in the kidneys and increase glucose excursion in the urine. Because their mechanism of action is independent of insulin secretion or action, they can be used in combination with other therapies [20]. T2D individuals treated with SGLT2i demonstrate improved glycemic control, increased glucose- and incretin-stimulated insulin secretion and enhanced insulin sensitivity as well as reduced blood pressure, decreased plasma lipids and reduced risk for cardiovascular events [21–24]. *Db/db* and *ob/ob* mouse models of obesity and T2D treated with SGLT2i (luseogliflozin, ipragliflozin or dapagliflozin) demonstrated reduced blood glucose, augmented insulin and GLP-1 secretion, increased β-cell mass, β-cell self-replication and α- to β-cell conversion, and restored β-cell identity [25–30]. Moreover, *db/db* mice with SGLT2 deletion showed improved insulin sensitivity and increased β-cell proliferation and decreased β-cell death [31]. Streptozotocin-induced type 1 diabetic mice treated with empagliflozin demonstrated increased insulin mRNA, serum insulin, and β-cell area and proliferation [32]. Although improved β-cell function by SGLT2i has been suggested in humans and rodents, the underlying mechanisms and timeframe of this effect remain elusive, with most studies performed in the setting of obesity and T2D.

Here we determined the underlying mechanisms and temporal progression to β-cell failure, and established the effects of SGLT2i therapy, in insulin secretory-deficient mouse model of K_ATP_-GOF induced NDM.

## 2. Materials and Methods

### Mouse models

Neonatal diabetic mice were generated as previously described [9]. Briefly, mice expressing Kir6.2 (K185Q,ΔN30) mutant transgene under the Rosa26 locus promoter were crossed with tamoxifen-inducible Pdx-Cre mice, to generate transgenic mice expressing the NDM mutation in β-cells. To induce transgene expression, 12 to14-week old mice were injected with five consecutive daily doses of tamoxifen (50 μg/g body weight). Littermate single transgenic (Rosa-kir6.2, Pdx-cre) were also injected with tamoxifen and used as controls since to date, no significant differences in phenotype were found. Both male and female mice were used for all experiments as no significant sex-based differences were found. Mice were maintained on a 12-hour light/dark cycle. Animals were house in BJCIH animal facility with a 24hs surveillance with veterinarian and personnel assigned to the facility. Power analysis determined the sample size, and it is stated in each experimental design and figures. Experimental groups were randomized, and treatment separation was assigned blinded.

### Ethics statement

This study was carried out in strict accordance with the recommendations in the Guide for the Care and Use of Laboratory Animals of the National Institutes of Health. The protocol was approved by the Washington University School of Medicine Animal Care and Use Committee (IACUC, Protocol Number: 19-1078). Tissue collection was performed at the end of the experiments at a predetermined time, with the animals euthanized by inhalation of Isoflurane in a chamber, and then cervical dislocation. All efforts were made to minimize suffering.

### Plasma hormones, and blood and urine glucose measurements

Blood glucose was measured either randomly or after an overnight fast as indicated by using the one-touch Bayer Contour T5 glucometer (Mishawaka, IN). Plasma insulin and glucagon levels were quantified using a rat/mouse insulin and mouse glucagon ELISA (Crystal Chem, IL). Urine glucose was analyzed by glucose-Autokit (Wako Diagnostics, Richmond, VA). Plasma lipids were measured by Washington University Diabetes Research Center Metabolic Tissue Function Core (http://diabetesresearchcenter.dom.wustl.edu/diabetes-models-phenotyping-core/). For all *in vitro* analyses, at least three independent animals were used. Cells and tissues from each animal were kept separated and analyzed individually. Data collection was stopped at predetermined, arbitrary time as 10 days after initiation of DAPA/vehicle treatment. No data were excluded. Further methods details are available in the following sections or in the Supplementary Materials section.

### Glucose tolerance tests (GTT) and Insulin tolerance tests (ITT)

GTTs and ITTs were performed after overnight or six-hour fast, respectively. Blood glucose was measured before (time 0) and after (15, 30, 45, 60, 90 and 120 min) intraperitoneal (ip) injection of 1.5 mg/kg dextrose (GTT) or 0.5 U/kg human insulin (ITT, HI-210 Lilly).

### Pancreatic Islet Isolation

For islet isolation euthanized mice were perfused through the bile duct with Hank’s solution containing collagenase type XI (Sigma, St. Louis, MO). Pancreases were removed, digested for 11 minutes at 37°C in water bath, hand shaken, and washed in cold Hank’s solution. Islets were handpicked under a stereo microscope in RPMI (11mM glucose) media (ThermoFisher Scientific) supplemented with fetal calf serum (10%), penicillin (100 U/ml), and streptomycin (100 μg/ml). For all assays involving islets, freshly isolated islets were immediately processed and freeze at −80C to avoid changes induced in the absence of DAPA.

### Insulin and proinsulin content measurement

Total insulin and proinsulin content were measured using rat/mouse insulin ELISA kit (Crystal Chem, IL) and proinsulin ELISA kit (Mercodia, Uppsala, Sweden) respectively from batches of ten size-matched islets after acid-ethanol extraction [12].

### Quantitative PCR analysis

Islet RNA extraction and qPCR analysis was performed as described previously [10]. Primers used to determine ER stress markers [33] and mature islet cell markers [10] have been published.

### Western blot analysis

15μg of total protein lysate was loaded per lane. Blots were incubated overnight with the following antibodies: Beta-actin (1:1000; EMD Millipore, MO), Proinsulin (1:1000; CST, MA), TXNIP (1:1000; MBL International, MA), sXBP1 (1:1000; Santa Cruz Biotechnology, TX), Prohormone convertase 1/3 (1:1000; CST, MA), Prohormone convertase 2 (1:1000; CST, MA), SERCA2b (1:1000, Santa Cruz Biotechnology, TX). Blots were washed and probed with RDye infrared fluorescent dye-labeled secondary antibody conjugates (1:10,000; LI-COR biotechnology). Fluorescence intensity was quantified by image studio Lite (LI-COR biotechnology).

### Immunohistochemical and morphometric analysis

Pancreases from control and NDM mice were fixed in 10% NBF, and paraffin-embedded after serial dehydration for sectioning. Four- to eight mice from each genotype were sampled on 5μm thick sections, 25μm apart (spanning the whole pancreas) and used for immunohistochemical/morphometric analysis. For morphometric analysis, at least 3 pancreatic sections from 3-8 mice from each genotype were covered systematically by accumulating images from non-overlapping fields on an inverted EXC-500 fluorescent microscope (Visual Dynamix, Chesterfield, MO). Hematoxylin-Eosin (HE) staining was carried out as described previously [10]. Briefly, slides were stained with guinea pig anti-insulin (Abcam, 1:100), mouse anti-glucagon (Cell signaling, 1:100), rabbit anti-somatostatin (Abcam, 1:500) antibodies and their distribution visualized using secondary antibodies conjugated with AlexaTM 488 or AlexaTM 594 (Molecular Probes, Eugene-OR) using an EXC-500 fluorescent microscope (Visual Dynamix, Chesterfield, MO). Whole pancreatic area and islet area were determined by H&E staining, and β-cell area and intensity by insulin staining (#C27C9 Rabbit mAb, Cell Signaling). Β-cell mass was calculated as the product of relative β-cell area to whole pancreatic area and pancreatic weight for each mouse.

### Reactive Oxygen Species

Islets were dispersed into single cells using Acutase (Sigma-Aldrich) and plated in RPMI complete media overnight on ploy-D-lysine (Sigma-Aldrich) coated glass-bottom culture dishes (MatTek Corp). After overnight culture, cells were incubated in KRB containing 2.8 mM glucose for one hour, followed by 16.7 mM glucose for one hour at 37°C. In the last 30 minutes of incubation, CellROX Deep Red Reagent (final concentration of 5μM, Molecular Probes ThermoFisher Scientific, MA) and live nuclear reagent (Hoeschst 33342, Molecular Probes) were added. After 30 min, cells were washed with PBS 3 times and imaged using a Leica DMI 4000B inverted microscope (Lecia microsystems, IL). Fluorescence intensity was quantified by image studio Lite (LI-COR biotechnology).

### Calcium measurements

The genetically encoded calcium sensor pRSET-RcaMP1h was a gift from Loren Looger (Addgene plasmid # 42874). The coding sequence of the RcaMP1h sensor was subcloned into the pShuttle vector, and recombinant adenovirus particles were produced following the AdEasy XL Adenoviral Vector System protocol (#240010 Agilent Technologies). Islets were transduced using a microfluidic device to obtain uniform infection of the islet cells throughout the islet volume and were incubated overnight before imaging. Imaging on islets was performed on the LSM880 inverted microscope (Zeiss Inc), using a heated stage-top incubator (Pecon GmbH) 24 hours post transduction. Islets were imaged in KRBH medium (NaCl 128.8 mM, KCl 4.8 mM, KH2PO4 1.2 mM, MgSO4 1.2 mM, CaCl2 2.5 mM, NaHCO3 5mM, HEPES 10 mM, pH 7.40, 0.1% BSA) supplemented with the desired glucose concentration. RcaMP1h fluorescence was excited with a 561 nm laser, and the emission was collected through a 570-650 nm bandpass.

### Dapagliflozin treatment

NDM, db/db and ob/ob mice were randomly separated into two treatment groups – vehicle or – Dapagliflozin (DAPA, SelleckChem, Houston, TX), a sodium-glucose transporter 2 inhibitor. DAPA was suspended in vehicle containing 30% Polyethylene glycol, 5% Propylene glycol, 0.4% tween-80 as recommended by the manufacturer. 12 to 14-week-old NDM mice were injected with 5 consecutive doses of tamoxifen to induce diabetes. Mouse groups were assigned randomly, and the study was not blinded. On day 7, NDM mice received a daily oral gavage of either vehicle or DAPA (10 mg/kg of body weight) for 10 days. 10-weeks old *db/db* and *ob/ob* mice also received a daily oral gavage of either vehicle or DAPA (10 mg/kg of body weight) for 10 days. Vehicle and DAPA treated mice were monitored through blood glucose measurements daily and blood serum collection and then euthanized for *ex vivo* analysis. Data collection was stopped at predetermined, arbitrary time as 10 days after initiation of DAPA/vehicle treatment. No data were excluded. Further methods details are available in the following sections or in the Supplementary Materials section.

### Statistical Analysis

Data are expressed as mean ± standard error of the mean. Statistical differences between two groups were determined using Student’s t-test and among several groups were tested using analysis of variance (ANOVA) with GraphPad PRISM version 8.0 (La Jolla, CA), assuming that the data is normally distributed for each variable. Significant differences among groups with *p<0.05, **p<0.01; ***p<0.001 and ****p<0.0001 are indicated in the Figures, and non-significant differences are not shown. The sample size, n, indicates the total number of biological samples.

## 3. Results

### Early loss of insulin content and proinsulin accumulation in islets from insulin secretory-deficient NDM mice

Upon induction of K_ATP_-GOF expression by tamoxifen, NDM mice developed severe diabetes and a marked decrease in plasma insulin levels (Fig. 1A,B) [9]. As predicted by the ‘switch off’ of glucose-stimulated insulin secretion, NDM mice showed significantly impaired glucose tolerance (Fig. 3C,D). Unexpectedly however, and not explained by expression of the K_ATP_ mutation, insulin content was markedly reduced (>50%) at day 15 of diabetes (Fig. 1E), correlating with decreased insulin immunostaining (Fig. 1F). This was accompanied by a significant increase in islet proinsulin protein levels (Fig. 1H), proinsulin:insulin content ratio (Fig. 1I) and plasma proinsulin (Fig. 1G). Prohormone convertases 1/3 and 2, which direct the endoproteolytic cleavage of proinsulin to insulin and C-peptide, were significantly decreased in NDM islets (Fig. 1J), suggesting reduced proinsulin processing as causal for proinsulin accumulation. As expected by the presence of overactive K_ATP_-channels in β-cells, NDM islets did not demonstrate an increase in cytosolic calcium in response to high glucose (Fig. 1K), confirming previous results in islets from long-standing diabetic NDM mice.

**Fig. 1.**
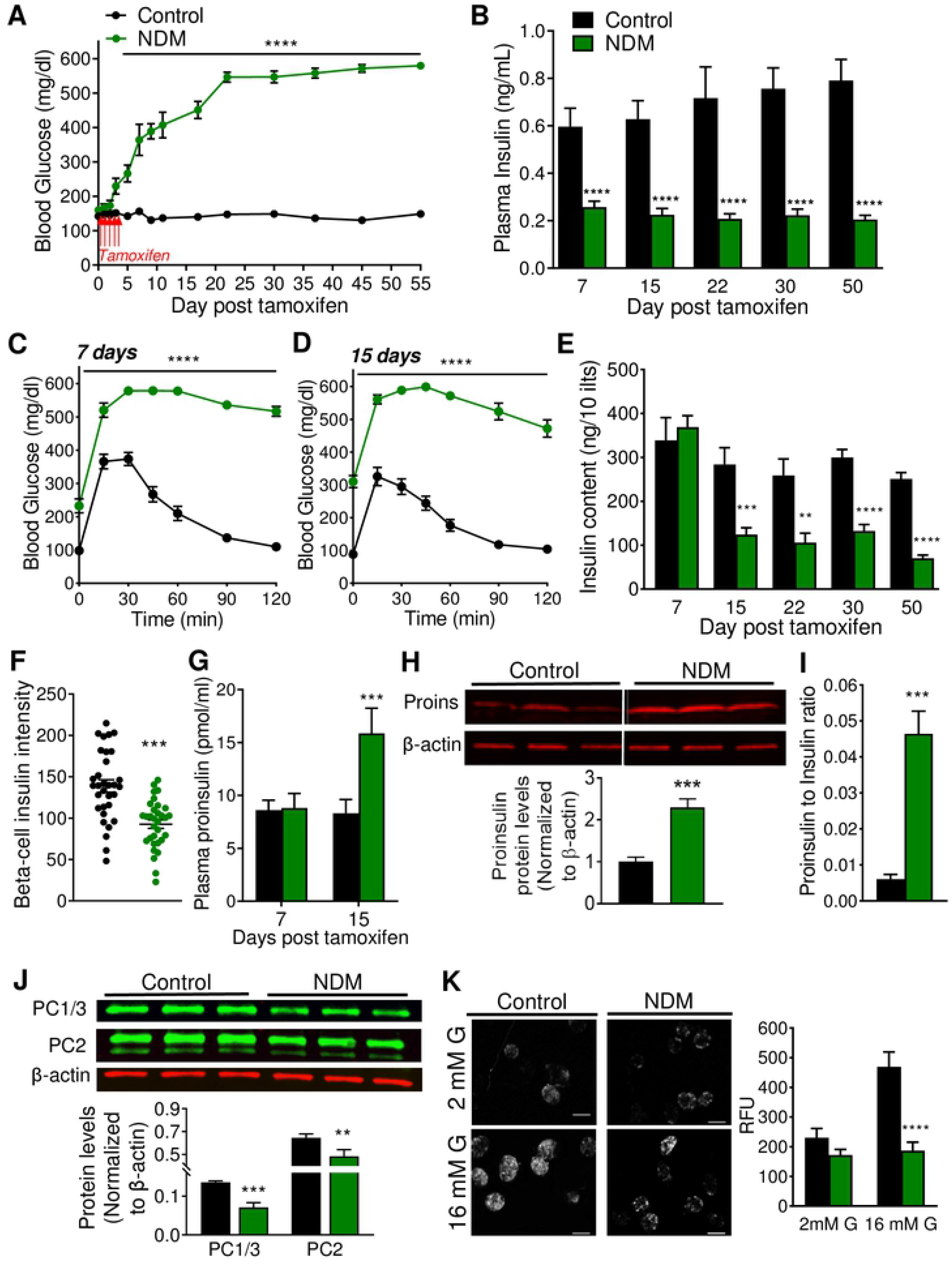
Loss of insulin content precedes β-cell mass and cell identity in NDM. (A) Non-fasting blood glucose and (B) plasma insulin over time (n=15-20 mice/group). Glucose tolerance test at day 7 (C) and 15 (D) post tamoxifen induction. (E) Total islet insulin content per 10 islets at different days post tamoxifen induction (n=8-10 mice/group). (F) beta-cell intensity determined in pancreatic sections immunostained with insulin. (G) Plasma proinsulin at day 15 post tamoxifen induction. (H) Western blot analysis of proinsulin protein, representative (top) and quantification (bottom) on islets at day 15 post tamoxifen induction, n=6 mice/group. (I) Proinsulin:insulin ratio at day 15 post tamoxifen induction (n= 10 mice/group) measured by Elisa. (J) Western blot analysis of prohormone convertases 1/3 and 2, representative (top) and quantification (bottom) on islets at day 15 post tamoxifen induction, n=4 mice/group. (K) Calcium imaging, representative left and quantification right at 2- and 16-mM glucose at day 15 post tamoxifen induction. Black=controls and green=NDM. Data are expressed as mean ± SEM. Significant differences \**P*<0.05, \*\**P*<0.01, \*\*\**P*<0.001, \*\*\*\**P*<0.0001.

### Increased oxidative and ER stress in islets from NDM diabetic mice

Proinsulin accumulation can activate the ER stress pathway response. Islets from NDM mice demonstrated a significant increase in mRNA levels in the ER stress markers *BiP*, spliced *XBP1* (*sXBP1)*. *ATF4* and *ATF6*, and no changes in *CHOP* (Fig. 2A). Protein levels of sXBP1, a product of the IRE pathway that is critical in highly secretory cells to maintain cell stress homeostasis, was significantly increased in NDM islets (Fig. 2B), and ER based ATPase sarco/endoplasmic reticulum calcium pump 2 (SERCA2b) significantly reduced (Fig. 2C). Reactive oxygen species (ROS) formation (Fig. 2D) and thioredoxin-interacting protein (TXNIP, a pro-oxidant protein) (Fig. 2E), were also markedly increased in NDM islets at day 15 of diabetes, suggesting increased oxidative stress in early stages of diabetes. Islets are particularly susceptible to oxidative damage due to low levels of antioxidants enzymes. Correlating with this, glutathione peroxidase and catalase were not detected, and superoxide dismutase 2 (SOD2) not altered in NDM islets (Fig. 2F). Interestingly, despite the decrease in insulin content at day 15 of diabetes, islet morphology, insulin and glucagon staining were not affected (Fig. 3A) in NDM islets. Moreover, total islet area, percent of β-cell area and β-cell mass (Fig. 3B). as well as the β-cell identity markers insulin, *Pdx1*, *Nkx6.1* and *Glut2* (Fig 3C) were not altered in NDM islets. In addition, plasma glucagon did not significantly change during diabetes progression (Fig. 3D), and percent of α-cell area and gene expression of the α-cell marker *Arx* remained similar in NDM and littermate controls islets at day 15 of diabetes (Fig. 3E,F).

**Fig. 2.**
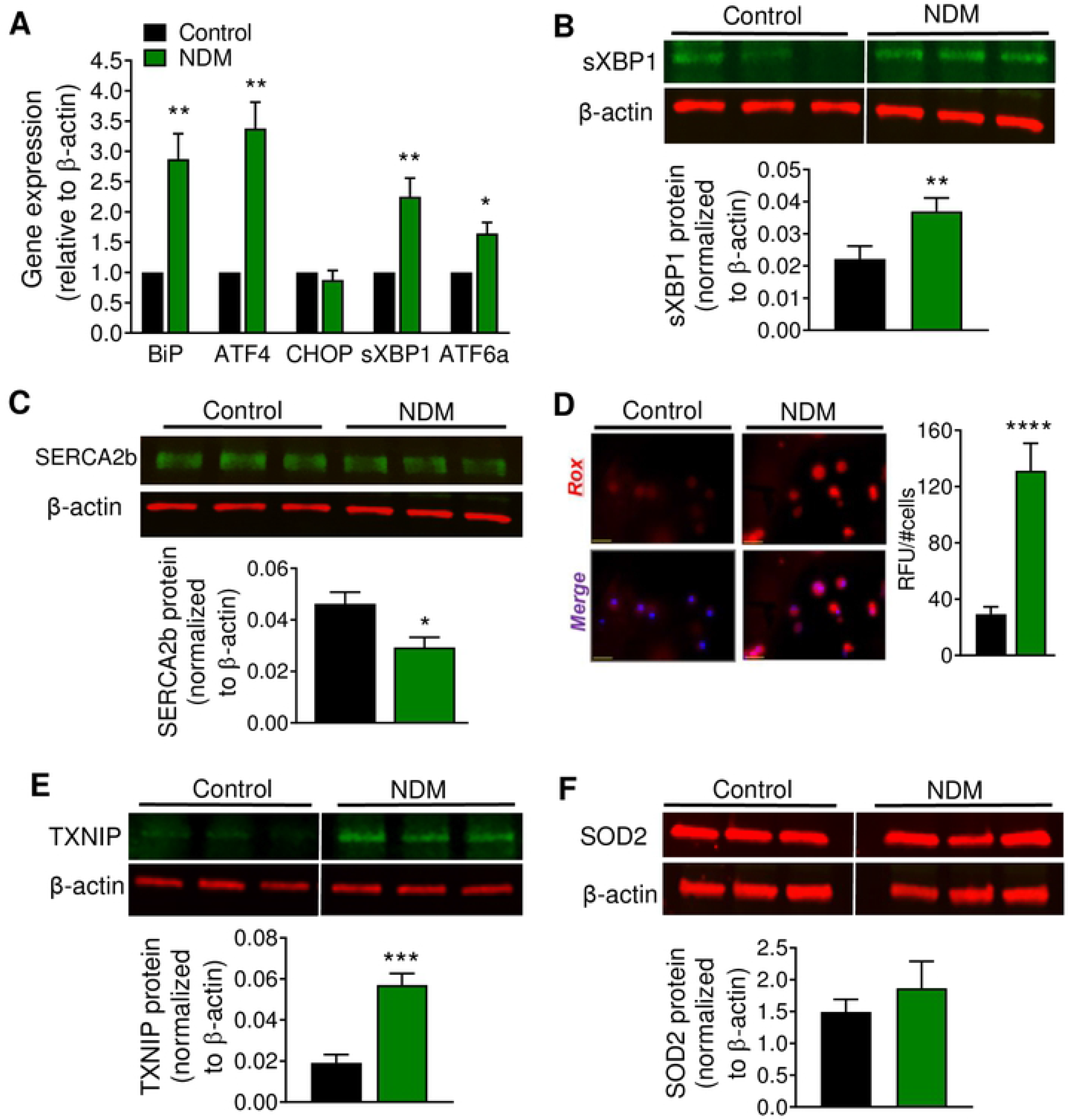
NDM islets demonstrated oxidative and ER stress in early diabetes. (A) Quantitative real-time PCR analysis of gene expression of ER stress markers in islets, n=5-8 mice/group. Western blot analysis of (B) sXBP1 and (C) SERCA2b, representative blot (top) and quantification (bottom), n=6 mice/group. (D) Reactive Oxygen Species (ROS) measurement in dispersed islets; representative images (left) and quantification (right), n=18 images from islets obtained from 3-4 mice in each group; scale bar=25 μm. (E) Representative western blot of TXNIP (top) and quantification (bottom), n=9 mice/group. Samples were run on the same gel but were not contiguous. (F) Representative western blot of SOD2 (top) and quantification (bottom), n=6 mice/group. Black=controls and green=NDM at day 15 post tamoxifen induction. Data are expressed as mean ± SEM. Significant differences \**P*<0.05, \*\**P*<0.01, \*\*\**P*<0.001, \*\*\*\**P*<0.0001.

**Fig. 3.**
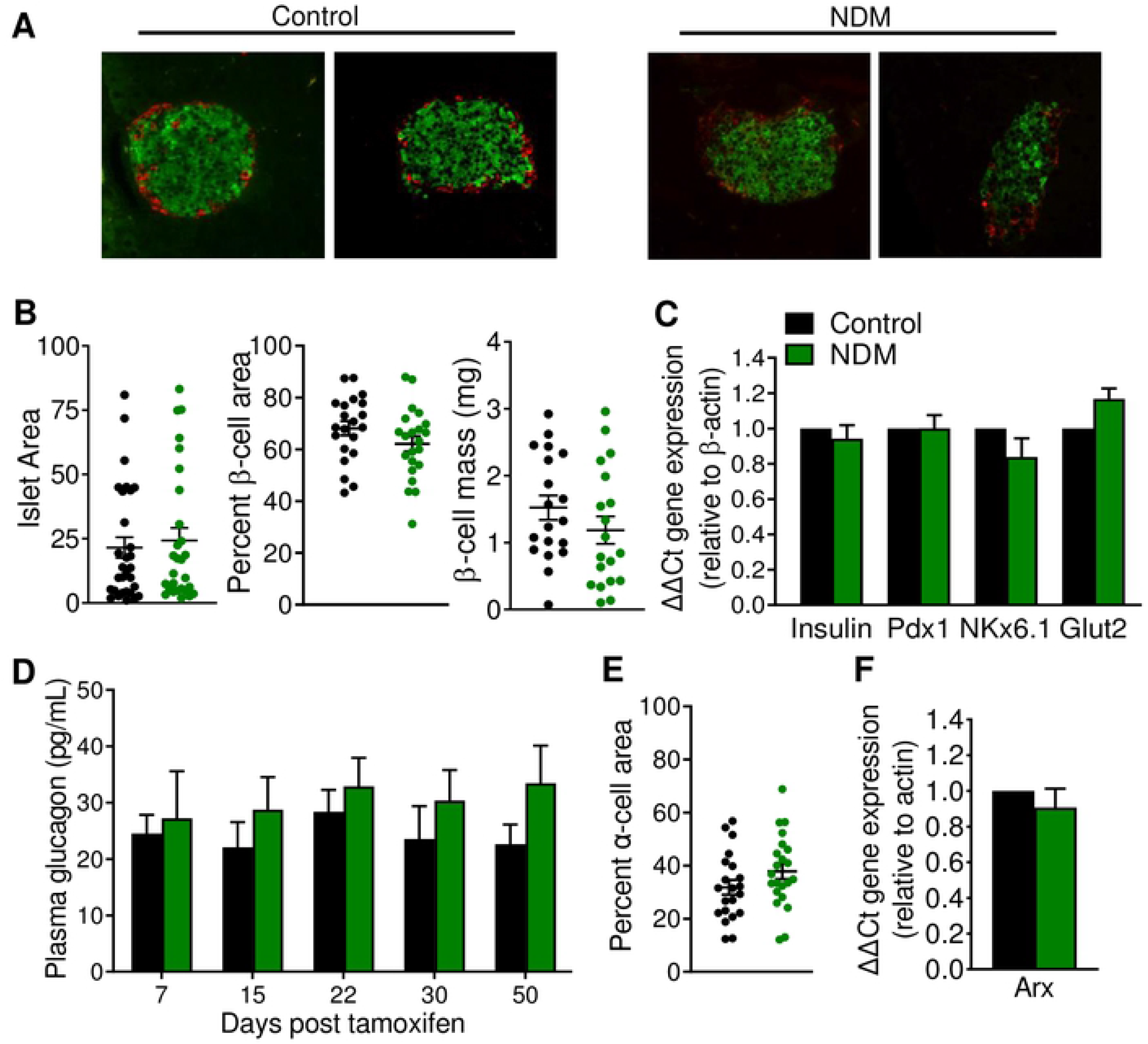
No changes in morphology or β-cell identity in islets from NDM mice at early stages of diabetes. (A) representative images of islets immunostained with insulin (green) and glucagon (red). (B) Islet area (left), percent of β-cell area (middle) and β-cell mass (right) n=8 mice/group, 3 slides from each mouse. (C) Quantitative real-time PCR analysis of gene expression of mature β-cell identity markers in islets (n=5-8 mice/group) at day 15 of diabetes. (D) Plasma glucagon overtime during development and progression of diabetes. (E) percent of α-cell area n=8 mice/group, 3 slides from each mouse, and (F) Quantitative real-time PCR analysis of gene expression of the α-cell identity marker *Arx* in islets, n=4 mice/group. Black=controls and green=NDM at day 15 post-tamoxifen injection. Data are expressed as mean ± SEM. Significant differences \**P*<0.05, \*\**P*<0.01, \*\*\**P*<0.001, \*\*\*\**P*<0.0001.

### Dapagliflozin therapy restores insulin content in non-obese NDM mice through reduction of β-cell stress

To determine whether reducing blood glucose alone reverts loss of insulin content and cellular stress, NDM mice were treated with the SGLT2i dapagliflozin (DAPA). As expected, DAPA-treated NDM mice showed a significant reduction in blood glucose (Fig. 4A), increased urinary glucose output (Fig. 4B), and no changes in plasma insulin (Fig. 4C) or body weight (Fig. 4D) after 10 days of DAPA therapy compared to vehicle-treated NDM. Plasma lipids such as triglycerides, cholesterol and free fatty acids (FFA) did not change by DAPA treatment (Fig. 4E). DAPA-treated NDM mice showed lower fasting glucose and improved glucose tolerance (Fig. 4F), and a mild, but not significant, improvement in insulin sensitivity (Fig. 4G) compared to vehicle-treated NDM mice. Ten days of DAPA treatment was sufficient to increase insulin content (Fig. 4H), reduce proinsulin (Fig. 4I) and proinsulin:insulin ratio (Fig. 4J) compared to vehicle-treated NDM mice. These changes were accompanied by a significant decrease in TXNIP and sXBP1 proteins (Fig. 4K), suggesting decreased cellular stress as a mechanism underlying recovery of β-cell function in NDM. No differences in any of the above-mentioned tests were observed in control mice treated with vehicle or DAPA (Fig. 4A-J). To test if alleviation of cellular stress is a unique effect of dapagliflozin, or an effect of lowering blood glucose, NDM mice were implanted with low dose slow-release insulin pellets (0.1U/day/pellet) 7 days post tamoxifen, to match the timeline for DAPA therapy (Fig. S1A). While blood glucose was significantly reduced, total islet insulin higher, and proinsulin:insulin ratio reduced in insulin-treated NDM mice compared to placebo treated mice (Fig. S1A-C), cellular stress marker proteins such as TXNIP and sXBP1 were not improved (Fig. S1D), suggesting unique effects of DAPA on reducing β-cell stress.

**Fig. 4.**
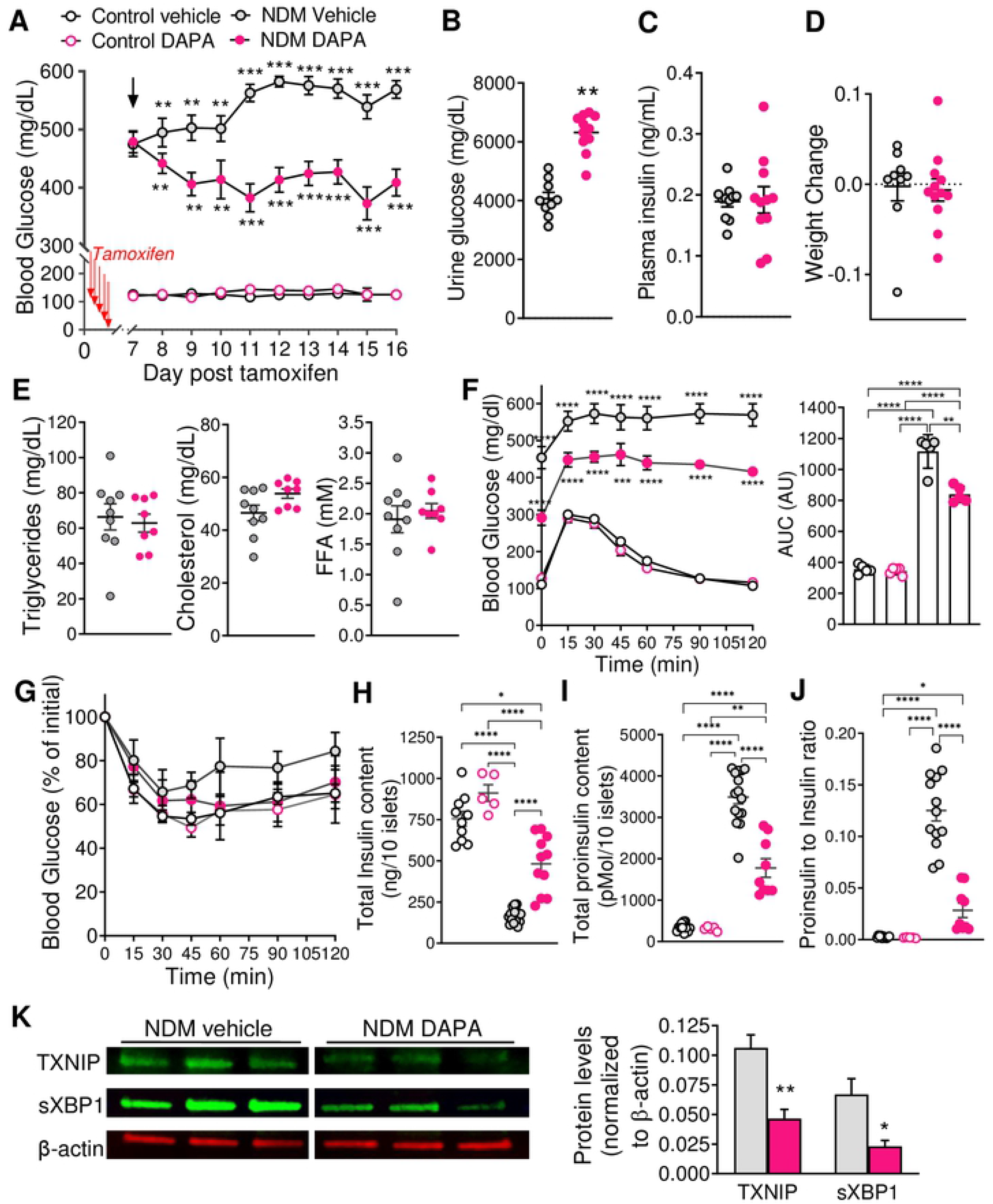
Dapagliflozin therapy in NDM mice improves β-cell function though reduction of oxidative and ER stress. (A) Blood glucose in control and NDM mice administered with vehicle or dapagliflozin (DAPA) by daily oral gavage on day 7 post-tamoxifen; n=5-15 mice/group. (B) Body weight (g) over time, (C) Plasma insulin and (D) urine glucose after 10 days of vehicle or DAPA treatment, n= 11 mice/group. (E) Lipid panel (tryglyceride, cholesterol and FFA) and (F) Glucose tolerance test (left) and calculated area under the curve (AUC, right) from control and NDM mice after 10-day treatment with vehicle or DAPA, n=5 mice/group. (G) Insulin tolerance test from control and NDM mice after 10-day treatment with vehicle or DAPA calculated as percentage from the initial blood glucose at 6-hrs fast (right), n=5 mice/group. (H) Islet total insulin content, (I) total proinsulin content, (J) proinsulin/insulin ratio and (K) prohormone convertases PC1/3 and PC2 from control and NDM mice after 10-day treatment with vehicle or DAPA, n=5-12 mice/group. (L) Western blot analysis on isolated islets after 10 days treatment with DAPA or vehicle. Representative blots (left) and quantification (right), n=4-6 mice/group. Black open circles: controls, pink open circles: control DAPA, grey filled circles: NDM vehicle treated, and pink filled circles: NDM DAPA treated mice. Arrow indicates initiation of DAPA therapy. Data are expressed as mean ± SEM. \**P*<0.05, \*\**P*<0.01, \*\*\**P*<0.001, \*\*\*\**P*<0.001.

Notably however, despite the significant decrease in blood glucose levels (Fig. 5A,B), insulin content (Fig. 5C) was not improved in islets from NDM mice when therapy was initiated at >42 days of diabetes. These results suggest inability of SGLT2i to improve β-cell function in animals with long-standing diabetes when they had already loss β-cell mass and identity.

**Fig. 5.**
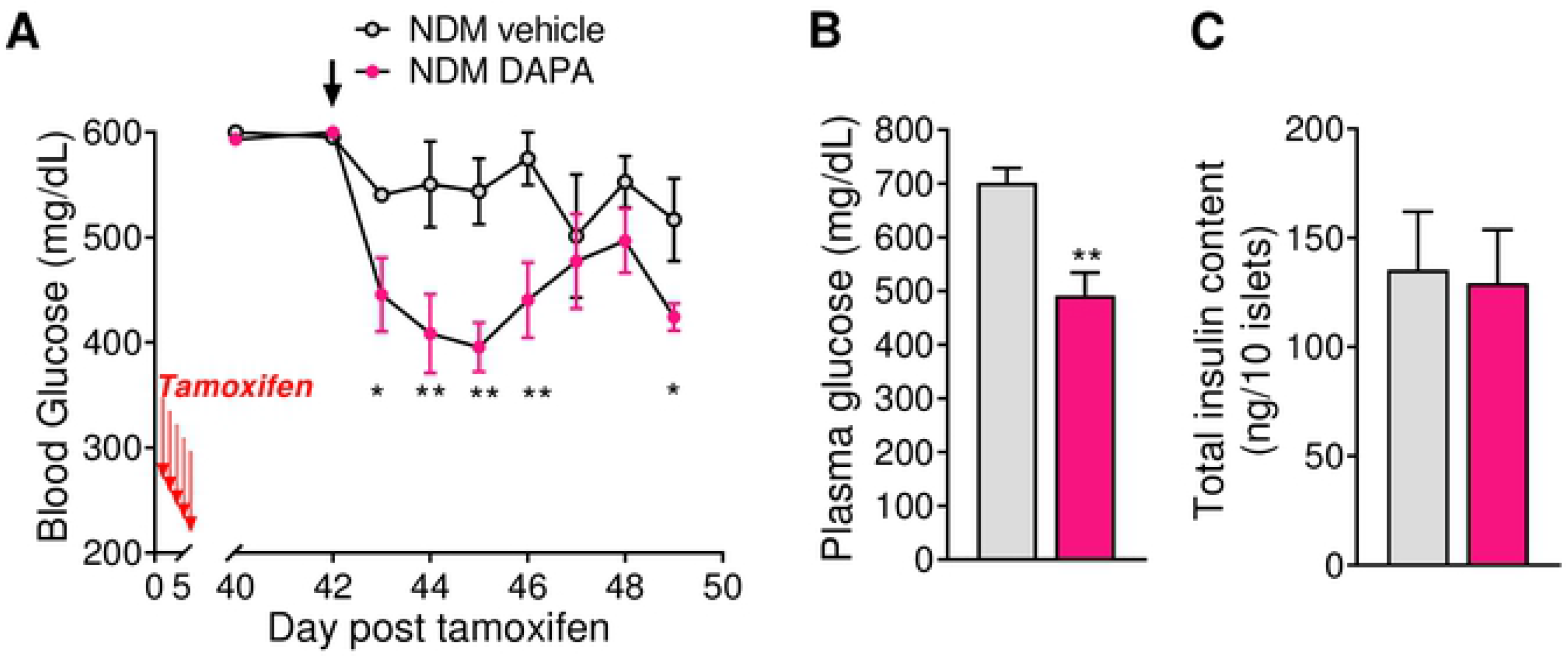
Dapagliflozin therapy does not improve insulin content in long-standing NDM. NDM mice treated with vehicle or DAPA, starting on day 42 post-tamoxifen (A) blood glucose over-time (left) and 10 days after treatment (right), n=6 mice/group. (B) Total islet insulin content 10 days after treatment with vehicle or DAPA, n=6 mice/group. Grey=vehicle-treated NDM and pink=DAPA-treated NDM mice. Arrow indicates initiation of DAPA therapy. Data are expressed as mean ± SEM. \**P*<0.05, \*\**P*<0.01.

### Dapagliflozin therapy reduced cellular stress in mouse models of obesity and T2D

Animal models of obesity and T2D diabetes such as leptin receptor deficient *db/db* and leptin deficient *ob/ob* mice were also treated with DAPA. DAPA-treated *db/db* mice showed a marked reduction in blood glucose (Fig. S2A), and no significant changes plasma insulin (Fig. S2B) or plasma lipids (Fig. S2C) compared to vehicle treated *db/db* mice. Similar to NDM mice, 10 days of DAPA therapy was sufficient to significantly increase insulin content (Fig. S2D) and markedly reduced TXNIP and sXBP1 protein levels (Fig. S2E). *Ob/ob* mice treated with DAPA also showed a reduction in blood glucose (Fig. S2F), although less consistent compared to the other two mouse models, and no significant changes in plasma insulin (Fig. S2G) or plasma triglycerides, cholesterol or FFA (Fig. S2H). DAPA-treated *ob/ob* mice also showed increased islet insulin content (Fig. S2I) and decreased sXBP1 and TXNIP (Fig. S2J).

## 4. Discussion

Loss of pancreatic β-cell mass and identity, and increased apoptosis have been demonstrated in T2D and T1D individuals [1, 3, 5, 34], and in animal models of monogenic diabetes as well as T1D and T2D diabetes [2, 4, 10, 11, 35]. Beta-cell oxidative and ER stress has been shown to be involved in β-cell failure in both T1D and T2D [34]. We demonstrated here that this is also the mechanism underlying early loss of insulin content in a mouse model of monogenic NDM, in the absence of hyperglycemia-induced insulin hypersecretion and β-cell exhaustion. Loss of insulin content, proinsulin accumulation and cellular stress are early events in NDM, leading to loss of β-cell mass and identity in long-standing diabetes. These results are in agreement with similar mechanism of β-cell failure occurring in human and mouse models of both T1D and T2D/obesity [34, 36, 37], thus highlighting common execution pathways for β-cell failure in different forms of diabetes.

Although some studies demonstrated that treatment with SGLT2i preserved β-cell mass in ZDF rats [38], streptozotocin-induced T1D mice [32] and in *db/db* and *ob/ob* mouse models of obesity and T2D [25–30, 39], the underlying mechanism of this protection remains elusive. We demonstrate here for the first time that SGLT2i therapy increased islet insulin content, and decreased proinsulin through reduction in β-cell oxidative and ER stress in mouse models of monogenic NDM and obese/T2D. Our demonstration of lack of restoration of insulin content in NDM mice treated with DAPA at day 42 of diabetes correlate with no improvements in insulin biosynthesis/secretion in 16 weeks old *db/db* mice treated with luseogliflozin [25], and with absence of preservation of β-cell function in 15-20 weeks old *db/db* mice treated with dapagliflozin [28], suggesting a point of no return for improvement of islet function by SGLT2i when β-cells had already lost their mature identity. These studies also correlate with those demonstrating that dapagliflozin was effective as monotherapy at early stages of human T2D, but only as add-on mediation to other antidiabetic drugs in individuals with long-standing diabetes [40].

In addition to the effect of lowering of blood glucose, SGLT2i also act as insulin-sensitizing agents. While some T2D individuals showed an improvement in insulin sensitivity, reduced HbA1c and weight loss by SGLT2i therapy [41], others did not [42]. Obese *db/db* mice treated with empagliflozin [39, 43], and *db/db* mice with genetic deletion of SGLT2 [31] demonstrated improved insulin sensitivity. However, lack of significant changes in insulin sensitivity and lipid profile in 10-day DAPA-treated NDM mice suggests additional effects of SGLT2i in insulin-resistant/obese states. Though, longer SGLT2i therapy could lead to reduction of plasma lipids and therefore additional improvements in β-cell function as it has been shown in mouse models of obesity/T2D [44–46].

Increased ER stress and decreased insulin signaling might synergistically reduce β-cell function [47–49], which is supported by our data demonstrating restoration of insulin signaling and reduction of ER stress by DAPA. Insulin therapy also preserved insulin content and decreased proinsulin:insulin ratio in NDM mice, which is consistent with proinsulin synthesis not regulated by insulin autocrine feedback on β-cells [49]. Therefore, we speculate that synergistic effects might be induced by a combination of SGLT2i and insulin therapy, which could explain the beneficial effects of combination of these two agents in individuals with long-standing T2D [40]. Thus, we may reconceptualize the role of SGLT2i as organ-protective agents in promoting adaptive cellular reprogramming of stressed cells for survival and function, as has been proposed elsewhere for cardiorenal benefits [50]. This paradigm shift in understanding additional actions of SGLT2i correlate with the reduction of β-cell stress in NDM mice by DAPA, but not by insulin therapy.

In conclusion, our study showed increased β-cell oxidative and ER stress as the underlying mechanism of early loss of insulin content in insulin secretory-deficient monogenic NDM, and that lowering blood glucose alone by SGLT2i is sufficient, through reduction of oxidative and ER stress, to restore insulin content and β-cell function. From a therapeutic perspective, our study strongly supports the use of SGLT2i as monotherapy in initial stages to preserve functional β-cell mass in monogenic diabetes, and as add-on combinational therapy to other antidiabetic agents in long-standing diabetes. Strikingly, similar underlying mechanisms for protection of β-cell failure by SGLT2i were found in different forms of diabetes suggesting common execution pathways.

## Abbreviations

NDM: neonatal diabetes mellitus
SGLT2: sodium glucose transporter 2
K_ATP_: ATP sensitive potassium channel
TXNIP: Thioredoxin Interacting protein
sXBP1: spliced X-box binding protein 1
ER: endoplasmic Reticulum
PC: proinsulin convertase
BiP: binding immunoglobulin protein
CHOP: C/EBP homologous protein
Pdx1: pancreas-duodenum homeobox 1
Nkx6.1: homeobox protein Nkx6.1
Glut2: glucose transporter 2
FFA: free fatty acid
DAPA: dapagliflozin
T2D: type 2 diabetes

## Author contributions

Z.A.S. and M.S.R designed the study and wrote the manuscript. Z.A.S, Z.Y. A.U., E.M.E. and M.S.R. performed experiments and data analysis. M.S.R conceptualized the study and edited the manuscript. All authors read and approved the final version of the manuscript.

